# Genomic Analyses of SLAMF7 CAR-T Cells Manufactured by *Sleeping Beauty* Transposon Gene Transfer for Immunotherapy of Multiple Myeloma

**DOI:** 10.1101/675009

**Authors:** Csaba Miskey, Maximilian Amberger, Michael Reiser, Sabrina Prommersberger, Julia Beckmann, Markus Machwirth, Hermann Einsele, Michael Hudecek, Halvard Bonig, Zoltán Ivics, on behalf of the CARAMBA consortium

## Abstract

Widespread treatment of human diseases with gene therapies necessitates the development of gene transfer vectors that integrate genetic information effectively, safely and economically. Accordingly, significant efforts have been devoted to engineer novel tools that i) achieve high-level stable gene transfer at low toxicity to the host cell; ii) induce low levels of genotoxicity and possess a ‘safe’ integration profile with a high proportion of integrations into safe genomic locations; and iii) are associated with acceptable cost per treatment and scalable/exportable vector production to serve large numbers of patients. The *Sleeping Beauty* (SB) transposon has been transformed into a vector system that is fulfilling these requirements.

In the CARAMBA project, we use SB transposition to genetically modify T cells with a chimeric antigen receptor (CAR) specific for the SLAMF7 antigen, that is uniformly and highly expressed on malignant plasma cells in multiple myeloma. We have demonstrated that SLAMF7 CAR-T cells confer specific and very potent anti-myeloma reactivity in pre-clinical models, and are therefore preparing a Phase I/IIa clinical trial of adoptive immunotherapy with autologous, patient-derived SLAMF7-CAR T cells in multiple myeloma (EudraCT Nr. 2019-001264-30/CARAMBA-1).

Here we report on the characterization of genomic safety attributes in SLAMF7 CAR-T cells that we prepared in three clinical-grade manufacturing campaigns under good manufacturing practice (GMP), using T cells that we obtained from three healthy donor volunteers. In the SLAMF7 CAR-T cell product, we determined the average transposon copy number, the genomic insertion profile, and presence of residual SB100X transposase. The data show that the SLAMF7 CAR transposon had been inserted into the T cell genome with the close-to-random distribution pattern that is typical for SB, and with an average transposon copy number ranging between 6 and 12 per T cell. No residual SB100X transposase could be detected by Western blotting in the infusion products. With these attributes, the SLAMF7 CAR-T products satisfy criteria set forth by competent regulatory authorities in order to justify administration of SLAMF7 CAR-T cells to humans in the context of a clinical trial. These data set the stage for the CARAMBA clinical trial, that will be the first in the European Union to use virus-free SB transposition for CAR-T engineering.

**Disclosures:** This project is receiving funding from the European Union’s Horizon 2020 research and innovation programme under grant agreement No 754658 (CARAMBA).

## INTRODUCTION

### *Sleeping Beauty* transposon system for nonviral engineering of CAR-T cells

The *Sleeping Beauty* (SB) transposon system [1] has been developed as a useful and promising tool for genetic engineering, including gene therapies (recently reviewed in [27]). The major advantage of SB gene delivery is that it combines the favorable features of viral vectors with those of naked DNA molecules, namely i) permanent insertion of transgene constructs into the genome by the transposition mechanism leads to sustained (potentially life-long) and efficient transgene expression in preclinical animal models (recently reviewed in [4-6]); ii) transposon vectors can be maintained and propagated as plasmid DNA, which makes vector manufacture scalable and cost-effective, and only requires S1 biological safety conditions, thereby significantly simplifying manufacture of the cell-based product. In addition, transposons can deliver larger genetic cargoes than viruses [8], and transposon-based vectors have an attractive safety profile [9-13].

One exciting area of therapeutic cell engineering with immediate translational relevance is adoptive immunotherapy with tumor-reactive T cells that are engineered by gene transfer to express a synthetic chimeric antigen receptor (CAR) [14, 15], resulting in selective tumor cell killing. The most advanced clinical development is the use of CARs specific for the B-lineage marker CD19. Several study groups have demonstrated that CD19 CAR-T cells are able to induce durable complete remissions in patients with chemotherapy- and radiotherapy-refractory B-cell acute lymphoblastic leukemia (ALL), non-Hodgkin lymphoma (NHL) and chronic lymphocytic leukemia (CLL) [16-23]. Two CAR-T cell products have obtained marketing authorization for the treatment of certain CD19-positive ALL and NHL.

SB has recently made a successful clinical debut in two studies of CD19 CAR-T cell therapy [24]. These studies showed that the administration of SB-engineered CD19-specific CAR-T cells is safe, and augments the Graft-versus-Leukemia effect in patients who had undergone prior hematopoietic stem cell transplantation. We have recently shown that CD19 CAR-T cells engineered by SB transposition and lentiviral transduction are equally effective in pre-clinical lymphoma models *in vitro* and *in vivo* [25], and that vectorization of the SB100X hyperactive transposase and a CAR transposon as minicircle (MC) DNA reduces toxicity and significantly improves transposition efficacy [25]. As a consequence, therapeutic doses of CD19 CAR-T cells can be obtained within as short as 14 days from 1 × 10^6^ input T cells with our MC-based SB approach, and the manufacturing process can be shortened to 72 hours or less, if higher input T cell numbers are utilized.

Another clinical proof-of-concept for CAR-T cell therapy has been obtained in multiple myeloma, a hematologic malignancy with clonal proliferation of plasma cells. The lead antigen for CAR-T cells in multiple myeloma is B cell maturation antigen (BCMA). A recent clinical trial with BCMA-specific CAR-T cells has highlighted their therapeutic potential with several partial and complete remissions [26]. However, downregulation of antigen expression and, as a consequence, the emergence of myeloma cell variants with antigen loss have been described [26], underscoring the need to identify and validate additional target antigens. We have recently shown that CAR-T cells targeting the alternative myeloma antigen SLAMF7 are effective in pre-clinical models [27]. We are preparing a multi-center clinical trial with SLAMF7-specific CAR-T cells engineered with SB gene transfer (http://www.caramba-cart.eu/).

## RESULTS

### Assessment of residual transposase as a safety parameter of a *Sleeping Beauty*engineered SLAMF7 CAR-T cell product

We applied a previously established protocol for *ex vivo* engineering of human T cells with SB vectors [25], albeit modified for GMP-compatibility and scaled to clinical production, stably delivering the SLAMF7 CAR by an electroporation procedure. In this procedure, SB100X transposase is encoded by mRNA and the SLAMF7 CAR transposon as MC. We generated three independent lots of SLAMF7 CAR-T cells from PBMCs of three independent healthy donor volunteers.

Stability of genomically integrated transgenes is of paramount importance for clinical applications. That is, ideally, transposition should take place only once during transfer of the therapeutic transgene from the transfected vectors into the cellular genome. Any further transposition event between chromosomal locations is undesired. One major determinant of potentially on-going transposition events in cell populations is the prolonged availability of the transposase. In this context, supplying the SB transposase in the form of mRNA has the added advantage of clearance of both the mRNA and the encoded transposase protein in the 14-day expansion period of CAR-T cell manufacturing. Importantly, the half-life of SB transposase has been estimated to be ∼80 h in cycloheximide-treated cultured cells [28].

In order to address the potential presence of the SB100X transposase at the end of the manufacturing period, cell populations from three independent validation runs were collected, and protein extracts analyzed by Western blotting alongside known amounts of recombinant purified SB100X transposase (**Fig. 1**). The detection limit of SB100X through chemiluminescent Western blotting in this procedure was 50 pg (**Fig. 1**), which corresponds to ∼7.66 × 10^8^ SB100X transposase molecules (SB100X = ∼39.29 kDa). Because we did not observe a band corresponding to the SB100X transposase in the SLAMF7 CAR-T cell extracts (**Fig. 1**), we conclude that the detection limit of this procedure is ∼766 SB100X molecules per cell, and all three validation runs contain residual SB transposase below this limit. Furthermore, taking into account the half-life of SB transposase protein [28] and the length of the culture process, we consider the SLAMF7 CAR-T cell product to be negative for residual transposase.

**Figure 1.**
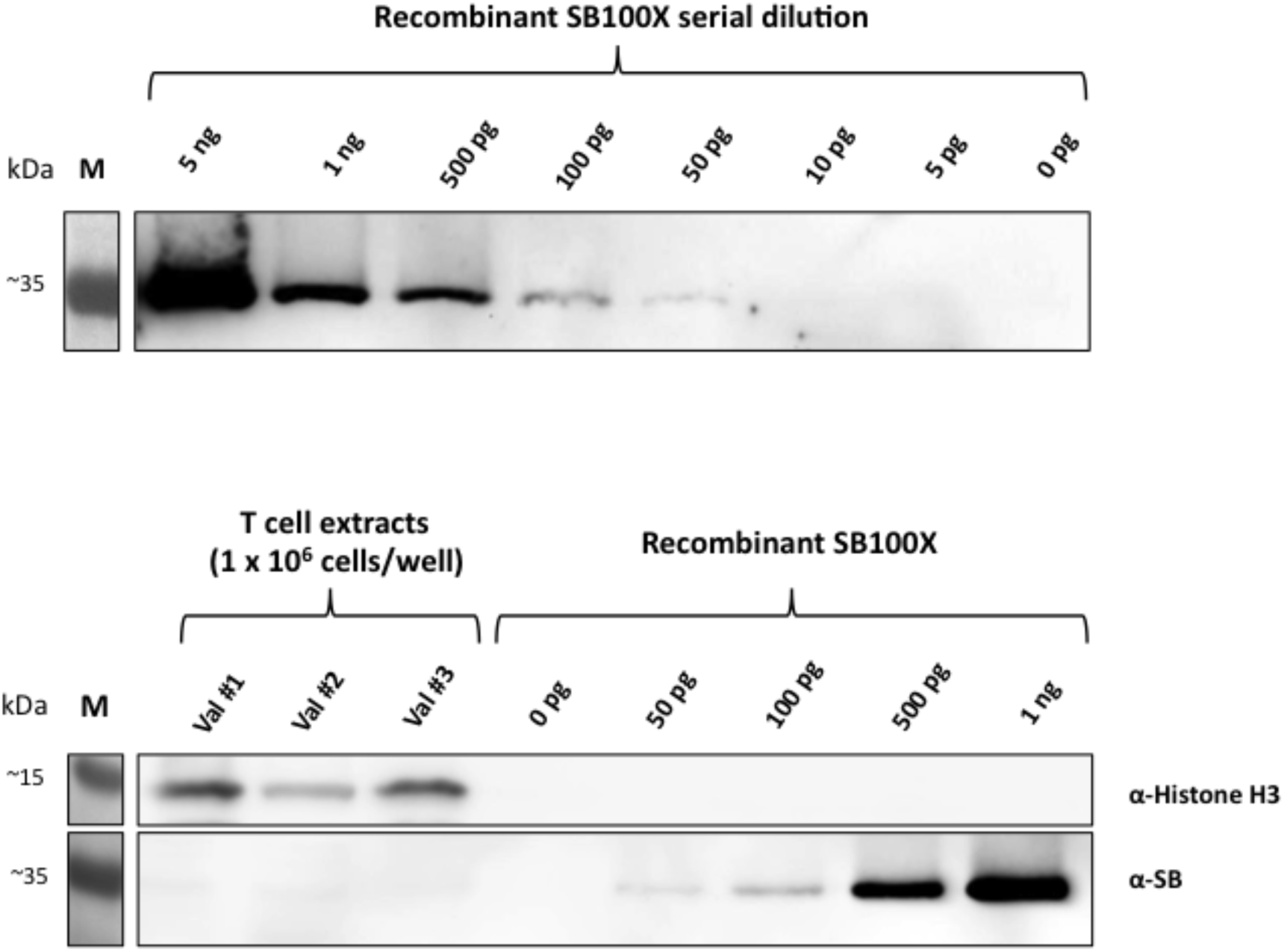
Western blotting establishes lack of detectable residual SB100X transposase in SLAMF7-positive CAR-T cell populations. A volume of cell extract corresponding to 1 × 10^6^ cells of each validation run was subjected to SDS-PAGE alongside recombinant SB100X protein in concentrations ranging from 0 pg – 1 ng and blotted onto a nitrocellulose membrane for subsequent chemiluminescent Western blotting. Exposure with a-Histone H3 antibody (loading control) was 30 sec, with a-SB antibody 20 min.

### Vector copy number of genomically integrated SLAMF7 CAR transposons

With any vector that integrates into chromosomes in a semi-random manner comes the potential risk of insertional mutagenesis leading to transcriptional activation or inactivation of cellular genes [29]. Such genotoxic effects can have devastating consequences for the cell and the whole organism, including the development of cancer. We consider at least two factors that contribute to the potential genotoxicity of an integrating vector system: vector copy number per genome and genome-wide integration profile.

We first determined the numbers of SB transposon integrations obtained per T cell genome. Quantitative droplet digital PCR (ddPCR) was performed using unselected bulk cell populations of SLAMF7 CAR-T cells harvested 14 days post-nucleofection. The average vector copy number (VCN) per diploid genome in the three validation runs was 8, 6 and 12 (**Fig. 2**), which falls within the range of VCNs typically seen with CAR-T cells manufactured with viral gene transfer systems. We nevertheless note that VCNs can potentially be decreased by applying limited amounts of transposon and transposase components [30, 31] at the electroporation step.

**Figure 2.**
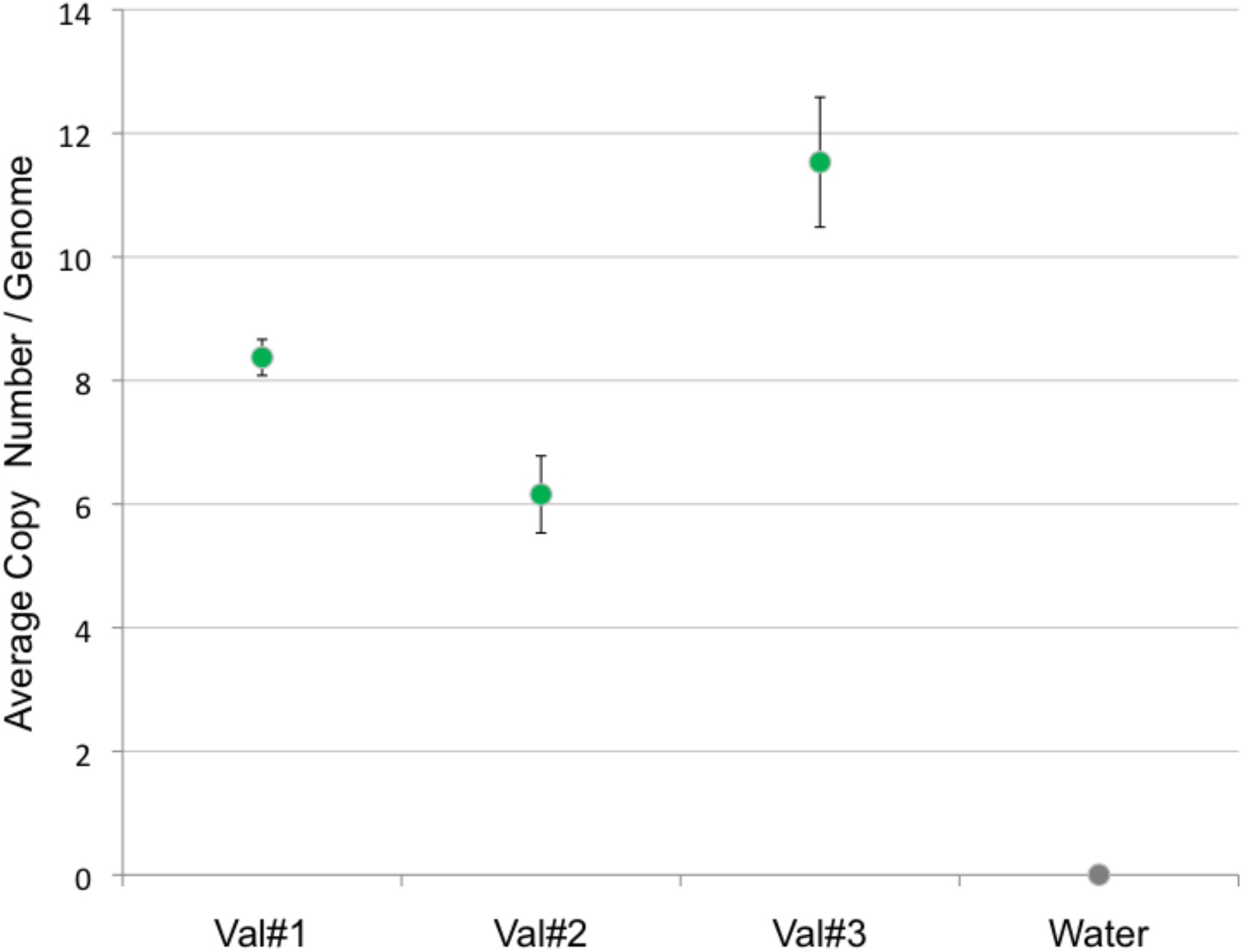
Vector copy numbers in bulk SLAMF7 CAR-T cells as determined by quantitative droplet digital PCR.

### Random insertion profile of the SLAMF7 CAR *Sleeping Beauty* vectors in the T cell genome

SB displays considerable specificity in target site selection at the primary DNA sequence level in that TA dinucleotides are near-obligate target sites [32]. We have previously undertaken a comparative study addressing target site selection properties of the SB and *piggyBac* (PB) transposons as well as MLV-derived gammaretroviral and HIV-derived lentiviral systems in primary human CD4^+^ T cells [33]. Our bioinformatic analyses included mapping against the T cell genome with respect to proximity to genes, transcriptional start sites (TSSs), CpG islands, DNaseI hypersensitive sites, chromatin marks and transcriptional status of genes. The SB transposon displayed the least deviation from random with respect to genome-wide distribution: no apparent bias was seen for either heterochromatin marks or euchromatin marks and only a weak correlation with transcriptional status of targeted genes was detected [33]. Our analyses collectively established a favorable integration profile of the SB transposon, suggesting that SB might be safer for therapeutic gene delivery than the integrating viral vectors that are currently used in clinical trials.

We constructed insertion site libraries from polyclonal SLAMF7 CAR-T cells for massive parallel sequencing on the Illumina MiSeq platform. We mapped and characterized from the three independent samples 5738, 6349 and 18574 unique insertion sites of the SLAMF7 CAR transposon. We detected the characteristic palindromic ATA**TA**TAT motif, which contains the TA dinucleotide target sequence of SB adjacent to all our MC-derived transposons (**Fig. 3**), similarly to what has been found for SB transposons mobilized from conventional donor plasmids [33, 34].

**Figure 3.**
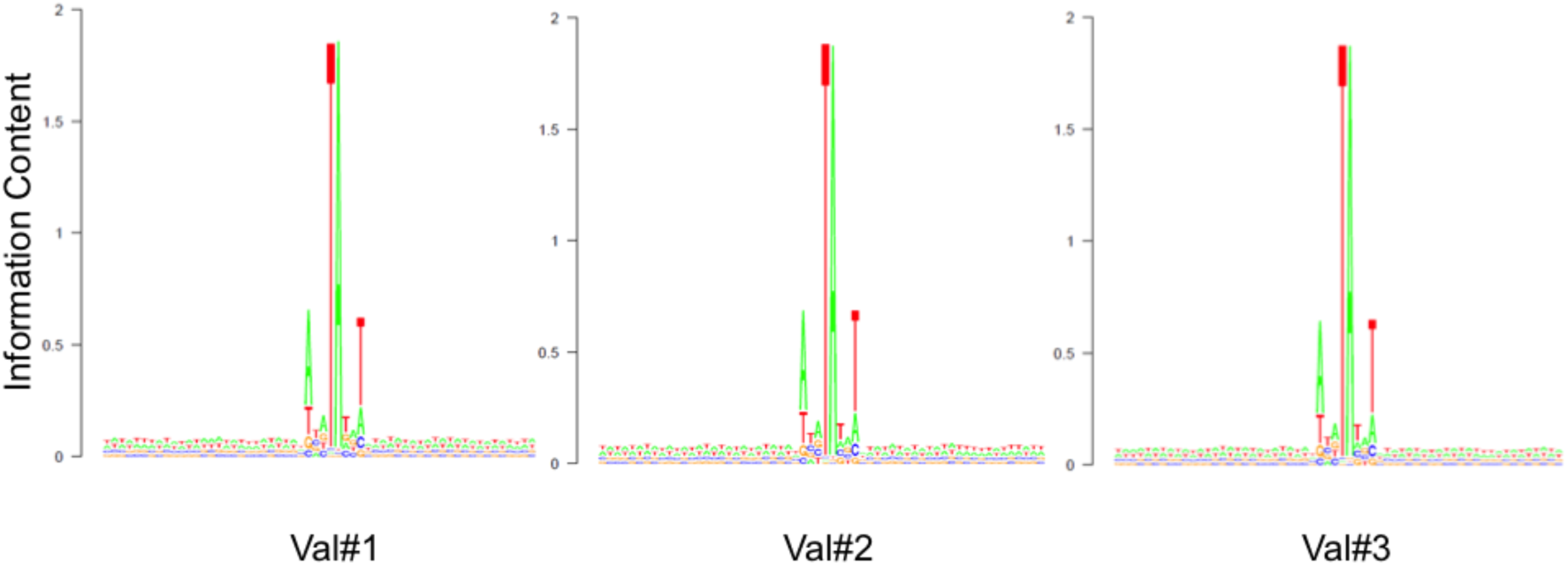
Canonical SB transposon integrations in the genomes of SLAMF7 CAR-T cells. The 60-nucleotide-long nucleotide frequency matrix was plotted using the SeqLogo tool in R software. The degree of base conservation is depicted by the height of the letters. Transposon integration occurs at the central TA dinucleotide of the consensus ATA**TA**TAT sequence.

We then analyzed whether there was a preference of SLAMF7 CAR transposon insertions into distinct sites of the genome, e.g. exons and introns, genes and cancerrelated genes. We found that transpositions had occurred with only a modest, yet statistically significant (p<0.001) bias towards genic categories (**Fig. 4**); however, in all evaluated categories this preference was substantially smaller than previously found for lentiviral and gammaretroviral integrations [25, 33]. Importantly, transposon insertions showed only a ∼1.2-fold enrichment in genes and a ∼1.4-fold enrichment in cancer-related genes relative to the expected random frequency. Concordantly, CAR transposons were also inserted into non-genic regions in a close-to-random manner (∼0.9-fold compared to random) (**Fig. 4**).

**Figure 4.**
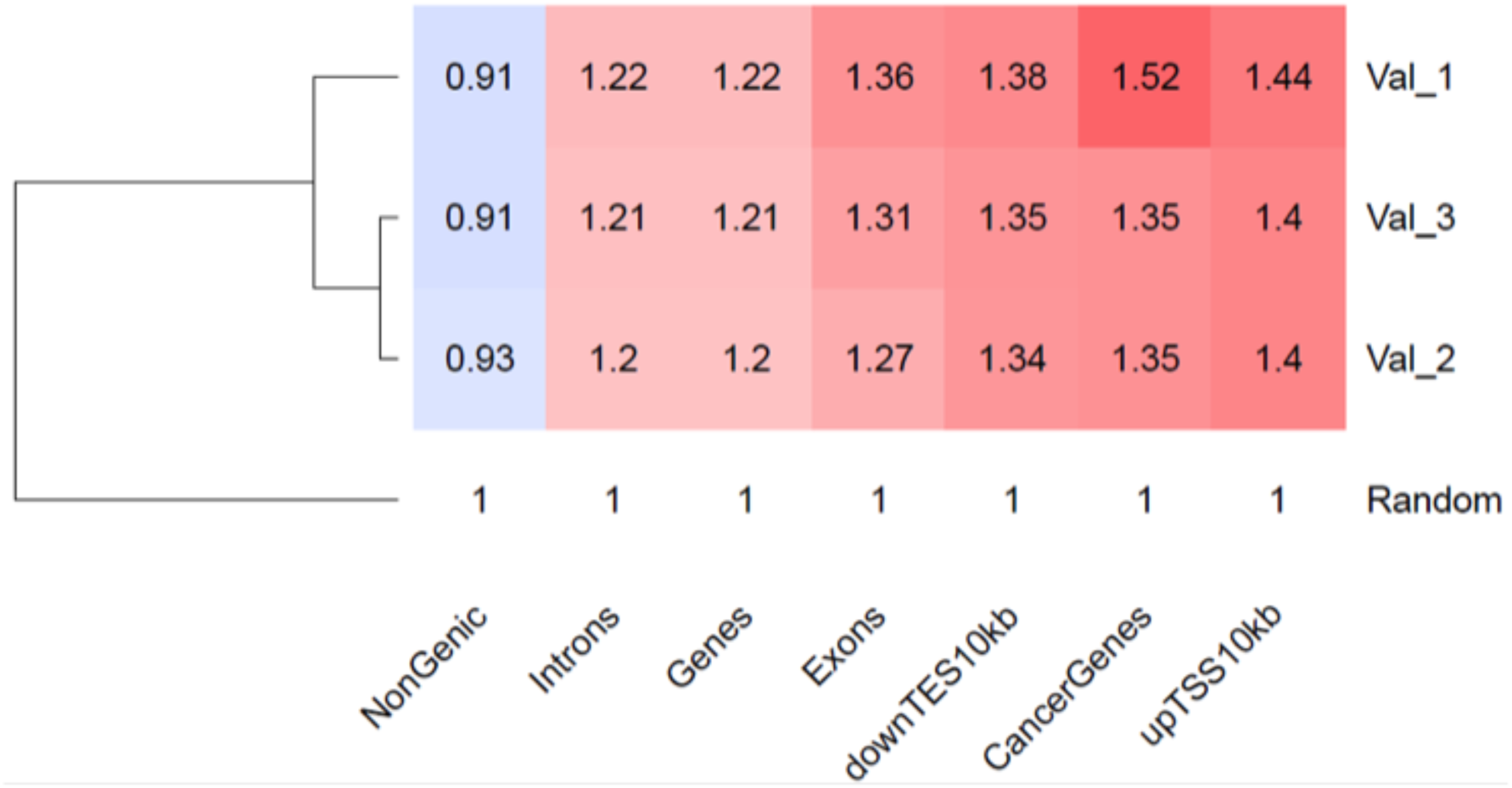
Genome-wide distribution of SLAMF7 CAR-containing SB transposons in the T cell genome. Distribution of SB insertions in functional genomic segments of human T cells. Numbers show relative enrichment above the random frequency (set to 1). Color intensities depict the degree of deviation from the expected random distribution (red: overrepresentation; blue: underrepresentation). *downTES10kb* stands for genomic regions extending 10 kb downstream from the transcriptional end sites of genes. *upTSS10kb* indicate 10-kb-long genomic segments upstream of transcriptional start sites of genes.

## DISCUSSION

Here we manufactured SLAMF7 CAR-T cells with a robust *ex vivo* protocol of virus-free gene delivery using the SB transposon system, and characterized these CAR-T cells with respect to three parameters that are important for the assessment of safety aspects associated with the engineered cell product: the presence of residual transposase at the end of the manufacturing cycle, the number of vector copies per target cell genome and genome-wide insertion profile of the integrated transgenes. Our results overall support the applicability of these cells for human therapy. First, we have not detected residual SB transposase in the CAR-T cells at day 14 post-electroporation. Because human cells do not express SB transposase endogenously, this finding suggests that transposition is confined to an initial time window after electroporation when transposition can occur from the transfected MCs into the T cell genome. At later timepoints, further rounds of transposition are inhibited in the absence of transposase, which is key to maintenance of genomic stability of the cell product. Second, average vector copy numbers of integrated SLAMF7 CAR transgenes per T cell genome fall within the range that is considered safe in the context of human T lymphocytes. Importantly, vector-driven insertional oncogenesis has not been observed to date in retroviral- and lentiviral-transduced terminally differentiated T cells [35]. Finally, random genomic distribution and lack of a pronounced preference for integration into genes (including oncogenes and tumor suppressors) further enhances the safety profile of SB gene delivery.

The advantages of using the SB system for gene therapy include the ease and reduced cost of manufacturing of clinical-grade, plasmid-based vectors compared to recombinant viral vectors [36], scalability of vector production, the ease of ensuring quality control for clinical use, and indefinite storage with absolute fidelity. In addition, there is pronounced flexibility with respect to a wide range of human cell types that are amenable to SB transposon-based nonviral gene delivery and with respect to a wide variety of transgene cassettes that can be efficiently incorporated into the target cell genome by this system. Indeed, it has recently been shown that allogeneic cytokine-induced killer (CIK) cells support highly efficient SB gene delivery, allowing the generation of CD19 CAR-CIK cells for clinical application in ALL patients [37]. In the context of adoptive T cell therapies, therapeutic T cell receptors (TCRs) can be efficiently introduced into T cell populations, and it was recently shown that the MC technology can boost TCR gene transfer with SB transposon vectors while reducing cellular toxicities [38]. Therefore, we consider SB transposition as a key enabling technology in order to make CAR-T cell therapy available on a global scale and in large patient cohorts.

## MATERIALS and METHODS

### Transposon vector components

The major functional components of MC-based transposon vectors have been previously described [25]. For the current work, the CD19-specific CAR was replaced by a SLAMF7-specific CAR fused with a T2A element and truncated epidermal growth factor receptor (EGFRt) [27]. A MC transposon vector encoding SLAMF7 CAR_EGFRt transposon was generated from the parental pT2-based plasmid by PlasmidFactory (Bielefeld) using site-specific recombination and purified by affinity chromatography. SB100X mRNA was produced by in vitro transcription by BioNTech (Idar-Oberstein).

### Generation of gene-modified T cells

CD8^+^ and CD4^+^ T cells were purified from PBMC and activated by stimulation with anti-CD3/anti-CD28 beads (Thermofisher) in separate cultures. Transfection of transposon and transposase vectors was performed on day 2 with 0,5 µg transposon MC and 2 µg SB100X transposase RNA per 1×10^6^ T cells on a 4D-Nucleofector according to the manufacturer’s instructions (Lonza). Transfected CD8^+^ and CD4^+^ T cells were expanded using the G-Rex culture system (Wilson-Wolf). On day 14, SLAMF7 CAR-expressing T cells were quantified using the EGFRt marker, and the SLAMF7 CAR-T product formulated to comprise CD8^+^CAR^+^ and CD4^+^CAR^+^ T cells at a 1:1 ratio.

### Western blotting

Cells were lysed in RIPA buffer (1 mM EDTA, 50 mM HEPES, pH 7.8, 150 mM NaCl, 1% (v/v) NP-40, 0.25% (v/v) Na-deoxycholate) with protease inhibitors at a concentration of 33.333 T cells/µl, 20 minute rotation at 4 °C followed by sonication (Bioruptor, Diagenode; two cycles on low power setting, 30 s ON, 30 s OFF). Clearing of the lysate was performed by centrifugation (15 minutes at 13000 rpm, 4 °C). A volume of cell extract corresponding to 1 × 10^6^ cells of each validation run was subjected to sodium dodecyl sulfate-polyacrylamide gel electrophoresis alongside recombinant SB100X (courtesy of C. Zuliani and O. Barabas, EMBL Heidelberg) protein in concentrations ranging from 0 pg – 1 ng and blotted onto a nitrocellulose membrane for subsequent Western blotting. SB100X was detected using a polyclonal SB transposase antibody (R&D Systems, AF2798) at a dilution of 1:1500 in blocking buffer (5% (w/v) powdered milk in TBS-Tween20 0.1% (v/v), pH 7.4), 1.5 h shaking at room temperature. For the loading control a polyclonal Histone-H3 antibody was used (Abcam, ab1791) at a dilution of 1:2000 in blocking buffer, shaking for 1.5 h at room temperature. Horseradish peroxidase conjugated secondary antibodies (Sigma Aldrich, A8919; Thermo Fisher Scientific, 31464) were used at dilutions of 1:5000 in blocking buffer shaking for 1 h at room temperature. Protein detection was performed using enhanced chemiluminescence detection (Amersham ECL Prime Western Blotting Detection Reagent).

### Vector copy number analysis

The procedure described in [31] was followed. 200 ng of genomic DNA (gDNA) was digested with 20-20 units of *Dpn*I and *Mlu*CI restriction enzymes (NEB) in 30 µl final reaction volume overnight in order to eliminate nonintegrated transposon MC DNA present in the samples and to fragment the gDNA, respectively. The samples were subjected to PCR amplifications using Taqman probes and primers specific for the transgene cargo of the transposon embracing three *Dpn*I recognition sites. Another set of primers and a Taqman probe specific for the single-copy human RPP30 gene to measure genome copy number of the pooled T cells. Triplicate PCR reactions were performed in 20 µl final volume with 10 ng of fragmented gDNA using the ddPCR Supermix for Probes (No dUTP) master mix (Biorad) with 11-11 pmol primers and 50-50 pmol of TaqMan probes (for sequences of primers and probes see **Table 1** below). The PCR droplets were generated in the QX100 device (Biorad). The program for the subsequent PCR was: 95 °C 10 min; 40 cycles of 94 °C 30 s, 50 °C 30 s, 60 °C 1 min; 98 °C 10 min. After thermal cycling the fluorescent droplets were counted in the QX100 Droplet Reader (Biorad) and genomic copy numbers were calculated with the Quanta Soft program (Biorad).

**Table 1.**
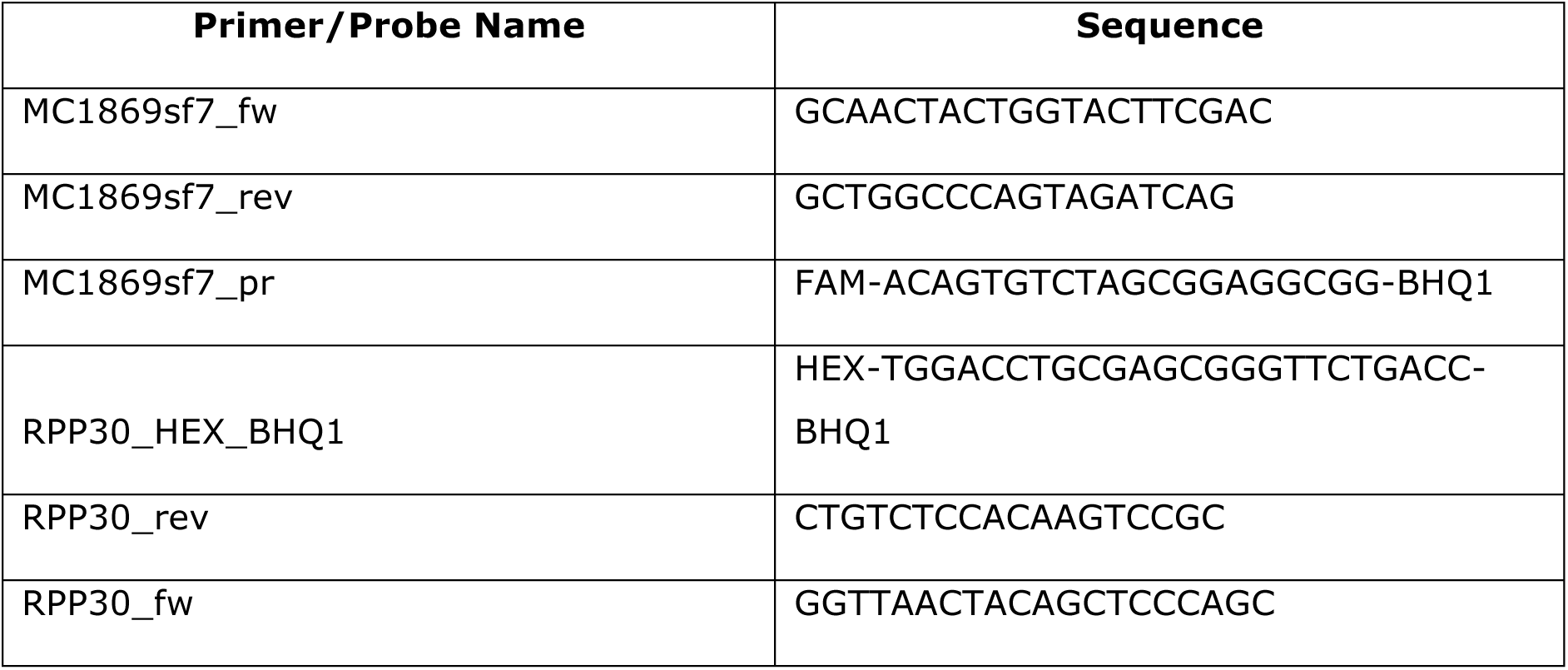
Sequences of primers used. BHQ = black hole quencher

### Insertion site analysis

The procedure described in [25] was followed. Genomic DNA of SLAMF7 CAR-T cells of three donors were isolated 14 days post-transfection. 2 µg DNA was sheared with a Covaris M220 ultra-solicitor device to an average fragment size of 600 bp in Screw-Cap microTUBEs in 50 µl, using the following settings: peak incident power 50 W, duty factor 20 %, cycles per burst 200, treatment 28 s. 1.2 µg of the sheared DNA was blunted and 5’-phosphorylated using the NEBNext End Repair Module (NEB), and 3’-A-tailed with NEBNext dA-Tailing Module (NEB) following the recommendations of the manufacturer. The DNA was purified with the Clean and Concentrator Kit (Zymo) and eluted in 8 µl 10 mM Tris pH 8.0 (EB) for ligation with 50 pmol of T-linker (see below) with T4 ligase (NEB) in 20 µl volume at 16 °C overnight. T-linkers were created by annealing 100 pmoles each of the oligonucleotides Linker_TruSeq_T+ and Linker_TruSeq_T-in 10 mM Tris-Cl pH 8.0, 50 mM NaCl, 0.5 mM EDTA. After heat-inactivation, ligation products enclosing fragments of non-integrated transposon donor plasmid DNA were digested with *Dpn*I (NEB) in 50 µl overnight, and the DNA was magnetic bead-purified and eluted in 20 µl EB. 5 µl eluate was used for the PCR I with 25 pmol of the primers specific for the linker and for the transposon inverted repeat: Linker and T-Bal-Long, respectively, with the conditions: 98 °C 30 s; 10 cycles of: 98 °C 10 s, 72 °C 30 s; 15 cycles of: 98 °C 10 s, ramp to 62 °C (1 °C/s) 30 s, 72 °C 30 s, 72 °C 5 min. All PCR reactions were performed with NEBNext High-Fidelity 2x PCR Master Mix (for PCR primer sequences see **Table 2** below). The PCR was bead-purified eluted in 20 µl EB and 5-5 µl of the elute were used for four parallel PCR II reactions with the primers: PE-nest-ind-N and SB-20-bc-ill-N for barcoding the samples of different T cell donors, using the following PCR program: 98 °C 30 s; 15 cycles of: 98 °C 10 s, ramp to 64 °C (1 °C/s) 30 s, 72 °C 30s, 72 °C 5 min. The final PCR products were separated on a 1 % agarose gel and the smears of 200-500 bp were gelisolated and purified. The libraries were sequenced on an Illumina MiSeq instrument using 2×300 sequencing setup.

**Table 2.**
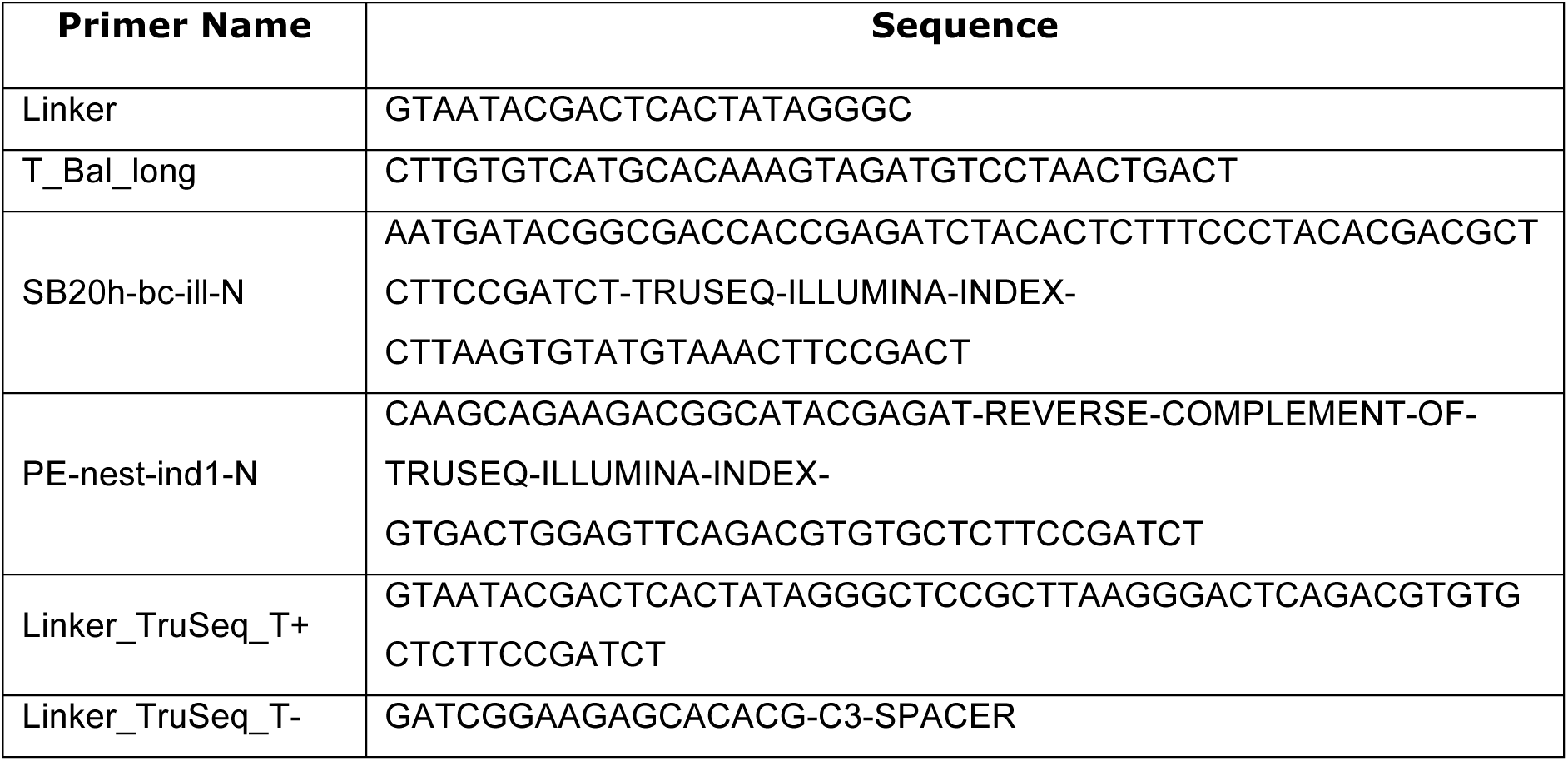
Sequences of primers used.

## Acknowledgments

We thank C. Zuliani and O. Barabas (EMBL Heidelberg) for providing purified SB100X transposase protein. This work was supported by funding from the European Union’s Horizon 2020 program for research and innovation under grant agreement No 754658 (CARAMBA). M. Hudecek is a member of the ‘Young Scholar Program’ (Junges Kolleg) of the Bavarian Academy of Sciences (Bayerische Akademie der Wissenschaften). Z. Ivics has been funded by the Deutsche Forschungsgemeinschaft (DFG) under grants IV 21/11-1 and IV 21/13-1, and has been supported by the Center for Cell and Gene Therapy of the LOEWE (Landes-Offensive zur Entwicklung Wissenschaftlich-ökonomischer Exzellenz) program in Hessen, Germany. H. Bonig has been supported by the Center for Cell and Gene Therapy of the LOEWE program in Hessen, Germany.

